# Altered phosphorylation of the proteasome subunit Rpt6 has minimal impact on synaptic plasticity and learning

**DOI:** 10.1101/719195

**Authors:** Samantha L. Scudder, Frankie R. Gonzales, Kristin K. Howell, Ivar S. Stein, Lara E. Dozier, Stephan G. Anagnostaras, Karen Zito, Gentry N. Patrick

**Author notes:** S.L.S. and F.R.G. contributed equally to this work. Author contributions: S.L.S., F.R.G., K.K.H., I.S.S., L.E.D., S.G.A., K.Z., and G.N.P. performed research; S.L.S., F.R.G., K.K.H., I.S.S., L.E.D., S.G.A., K.Z., and G.N.P. analyzed data; S.L.S., F.R.G., and G.N.P. wrote the paper. The authors declare no competing financial interests. To whom correspondence should be addressed: Gentry N. Patrick, Section of Neurobiology, Division of Biological Sciences, University of California San Diego, 9500 Gilman Drive, La Jolla, CA 92093-0366.

## Abstract

Dynamic control of protein degradation via the ubiquitin proteasome system is thought to play a crucial role in neuronal function and synaptic plasticity. The proteasome subunit Rpt6, an AAA ATPase subunit of the 19S regulatory particle, has emerged as an important site for regulation of 26S proteasome function in neurons. Phosphorylation of Rpt6 on serine 120 (S120) can stimulate the catalytic rate of substrate degradation by the 26S proteasome and this site is targeted by the plasticity-related kinase calcium/calmodulin-dependent kinase II (CaMKII), making it an attractive candidate for regulation of proteasome function in neurons. Several in vitro studies have shown that altered Rpt6 S120 phosphorylation can affect the structure and function of synapses. To evaluate the importance of Rpt6 S120 phosphorylation in vivo, we created two mouse models which feature mutations at S120 that block or mimic phosphorylation at this site. We find that peptidase and ATPase activities are upregulated in the phospho-mimetic mutant and downregulated in the phospho-dead mutant (S120 mutated to aspartic acid (S120D) or alanine (S120A), respectively). Surprisingly, these mutations had no effect on basal synaptic transmission, long-term potentiation, and dendritic spine dynamics and density in the hippocampus. Furthermore, these mutants displayed no deficits in cued and contextual fear memory. Thus, in a mouse model that blocks or mimics phosphorylation at this site, either compensatory mechanisms negate these effects, or small variations in proteasome activity are not enough to induce significant changes in synaptic structure, plasticity, or behavior.

## INTRODUCTION

Synaptic plasticity, which is fundamental for learning and memory, is mediated by both morphological and functional modifications to synapses. Furthermore, there is a growing body of evidence which suggests that impairments in synaptic plasticity contribute to neuropsychiatric disorders and neurodegenerative diseases. Emerging evidence has shown that local protein synthesis and degradation are crucial for the remodeling of synapses (Alvarez-Castelao & Schuman, 2015). Protein homeostasis, the balance between protein synthesis and degradation, is critical for maintaining cell health and viability. The Ubiquitin Proteasome System (UPS) is a major pathway through which proteins are broken down and recycled in eukaryotic cells, making it pivotal in their development, maintenance, and survival. The 26S proteasome is a multi-subunit complex which breaks down proteins targeted for degradation (Hershko & Ciechanover, 1998), and consists of a 20S catalytic core and a 19S regulatory particle (RP). The 19S regulatory particle includes a six subunit hexameric ring of AAA ATPases (Rpt1-6) which serve as the interface with the alpha subunits of the core particle (CP) and which provide the mechanical energy necessary to deubiquitinate substrates and move them into the catalytic core (Lander *et al.*, 2012; Matyskiela *et al.*, 2013). Protein homeostasis is particularly important in the CNS, as rapid response to external cues, maintaining and remodeling synaptic connections, and constantly adjusting the protein composition of pre- and postsynaptic compartments is critical in synaptic function and dysfunction (Ding *et al.*, 2006; Graham & Liu, 2017).

In mammals, studies have shown that proteasome inhibition alters basal synaptic transmission, long-term potentiation (LTP) and long-term depression (LTD) (Fonseca et al., 2006; Hou et al., 2006; Karpova et al., 2006; Deng et at., 2007). Interestingly, there appears to be a strict balance between new protein synthesis and protein degradation via the proteasome. While the application of protein synthesis inhibitors or proteasome inhibitors alone block LTP, the application of these drugs together occludes the effect of each other as LTP is unaltered [Fonseca et al. 2006 Neuron]. These studies highlight the idea that the relative concentration of key synaptic proteins can be rate-limiting for long-lasting changes in synaptic efficacy. We have previously shown that the blockade of action potentials with tetrodotoxin or up-regulation of activity with bicuculline inhibit and increase proteasome peptidase activity, respectively (Djakovic et al., 2009). Ca^2+^/calmodulin-dependent protein kinase II alpha (CaMKIIα) has been directly linked to this phenomenon, as it was demonstrated that CaMKIIα acts as a scaffold protein necessary for proteasome translocation into spines and also stimulates proteasome peptidase activity by phosphorylating serine 120 (S120) on Rpt6, an ATPase subunit on the 19S regulatory particle of the proteasome, in an activity-dependent manner (Bingol & Schuman, 2006; Djakovic *et al.*, 2009; Bingol *et al.*, 2010; Djakovic *et al.*, 2012). In subsequent *in vitro* investigations, we found that homeostatic scaling of synaptic strength induced by chronic changes in synaptic activity is impaired by altered Rpt6 S120 phosphorylation (Djakovic *et al.*, 2012). It was also demonstrated that the acute proteasomal inhibition or ectopic expression of an Rpt6 S120A phospho-dead mutant blocks activity-dependent new spine generation (Hamilton *et al.*, 2012). Furthermore, phosphorylation of S120 is increased during fear conditioning, and CaMKIIα-dependent regulation of proteasome phosphorylation and activity promotes memory consolidation (Jarome et al., 2013; Jarome et al., 2016). Together, these data suggest that CaMKIIα-dependent phosphorylation of Rpt6 at S120 may be an important regulatory mechanism for proteasome-dependent control of synaptic remodeling in various plasticity and behavioral paradigms.

We sought to further investigate the importance of Rpt6 phosphorylation and proteasome function in neurons. Historically, proteasome inhibitors have been used to understand the role of the UPS in synaptic plasticity and behavior. However, they are quite toxic to cells and therefore have a limited utility. Many loss of function mutations in proteasome subunits cause lethality in various organisms, therefore the need for more subtle mutant models which partially alter proteasome function is quite high. Here, we have generated two novel knock-in mouse models which eliminate Rpt6 S120 phosphorylation dynamics by mutating Rpt6 S120 to either alanine (S120A), which blocks phosphorylation, or to aspartate (S120D), which mimics phosphorylation due to its negatively charged carboxyl group. With these mouse models, we set out to examine the effects of altered Rpt6 S120 phosphorylation in the brain.

We found that affinity-purified proteasomes from Rpt6 S120A and S120D mutant mice exhibit alterations in kinetic rate of ATP hydrolysis and substrate degradation when compared to wild type littermates. Interestingly, however, we did not observe any significant changes in basal synaptic transmission and hippocampal long-term potentiation (LTP). In addition, spine density and outgrowth dynamics were unaltered in Rpt6 S120A mutants. Furthermore, fear memory, locomotor behavior, and anxiety-related behaviors were normal in both Rpt6 S120A and S120D mutants. These findings are in stark contrast to previous data which showed significant differences in synapse strength, synaptic scaling, and activity-dependent spine outgrowth (Djakovic *et al.*, 2009; Djakovic *et al.*, 2012; Hamilton *et al.*, 2012). This suggests that in neurons additional mechanisms may exist to compensate for alterations in proteasome function and mediate the tight control of the synaptic proteome required for proper changes in synaptic structure, function, and behavior.

## MATERIALS AND METHODS

### Generation of Rpt6 S120A and S120D knock-in mice

We generated Rpt6 phospho-mimetic (ser120 to aspartic acid; S120D), and phospho-dead (ser120 to alanine; S120A) KI mice (ingenious Targeting Laboratory; www.genetargeting.com). The targeting vectors were linearized and transfected by electroporation into BA1 (C57Bl/6 x 129/SvEv) (Hybrid) embryonic stem cells. Selected clones were expanded for southern blot analysis to identify recombinant ES clones (data not shown). The ES clones were microinjected into C57BL/6 blastocysts. After germline transmission, the Neo cassette was removed by mating to C57BL/6 FLP mice. Tail DNA was analyzed by PCR to identify heterozygous mice and verify deletion of the Neo cassette. Mutant heterozygous mice were backcrossed to C57BL/6. By visual inspection, Rpt6 S120D and S120A homozygous mutants (confirmed by PCR and sequencing) obtained by crossing heterozygous mutants, displayed normal body size, feeding, and mating behaviors. The intercross of heterozygotes resulted in production of wild-type, heterozygous, and homozygous offspring at the expected 1:2:1 Mendelian ratio. All procedures were approved by the UCSD IACUC and compliant with the NRC Guide.

### Antibodies and reagents

Anti-20S core α subunits monoclonal mouse antibody (mAb MCP231) and Rpt6 regulatory subunit monoclonal mouse antibody (mAb p45-110) were purchased from Enzo Life Sciences. Custom rabbit (pAb; Clone #07) anti-Rpt6 phospho-specific antibody for serine 120 (pS120) was previously generated commercially (ProSci) against a synthetic phosphorylated peptide (Djakovic *et al.*, 2012). N-succinyl-Leu-Leu-Val-Tyr-7-amino-4-methylcoumarin (Suc-LLVY-AMC) substrate was used (BACHEM). GST-Ubl and His-UIM constructs were a gift from the Alfred Goldberg lab. Malachite Green and Adenosine triphosphate were purchased from Sigma Aldrich. PhosSTOP kinase/phosphatase inhibitor and cOmplete protease inhibitors were purchased from Roche. Picrotoxin was purchased from Tocris.

### Proteasome purification and Mass Spec Analysis

Purification was conducted as previously described (Besche & Goldberg, 2012), with some modifications. Whole brain lysates were dounce homogenized in affinity purification buffer (APB: 25 mM HEPES–KOH, pH 7.4, 10% glycerol, 5 mM MgCl_2_, 1 mM ATP, and 1 mM DTT) and lysates cleared by ultra-centrifugation at 100,000g for 60 minutes. Lysate was incubated with purified GST-Ubl recombinant protein at a concentration of 0.2 mg/ml, then GSH-agarose beads were added and incubated for 2 hours at 4°C. The slurry was loaded onto a 20 ml column and washed twice with 10 ml APB. Proteasomes were eluted with purified UIM (2 mg/ml) in two incubations of 250 µl each. The resulting purified proteasomes were then measured for protein concentration, aliquoted, and frozen at −80°C. Trypsinized samples were directly subjected to mass spectrometry analysis.

### Peptidase and ATPase assays

Peptidase assays were conducted according to methods previously described (Kisselev & Goldberg, 2005). For peptidase assays, purified proteasomes were mixed with assay buffer (50 mM HEPES (pH 7.8), 10 mM NaCl, 1.5 mM MgCl_2_, 1 mM EDTA, 1 mM EGTA, 250 mM sucrose, 5 mM DTT, 2 mM ATP) and suc-LLVY-AMC substrate, 100 µl was loaded into a Costar black 96 well microassay plate (in triplicate), and the kinetic rate of cleavage was monitored by the increase in fluorescence (Excitation: 360 nm; Emission: 465 nm) at 37°C with a microplate fluorimeter (Perkin-Elmer HTS7000 Plus) every 30 seconds for two hours. The slope of the plot was calculated as the relative kinetic rate of substrate cleavage by the proteasome. The Malachite green assay, a colorimetric assay used to measure the evolution of inorganic phosphate from a mixture of pure proteasomes and ATP, was adapted from previous studies (Lanzetta *et al.*, 1979) to quantify ATP hydrolysis. The assay was conducted in triplicate on clear Costar 96 well plates, and the absorption at 660 nm was measured via plate reader spectrophotometer at different time points. The slope of the graph was calculated as the kinetic rate of ATP hydrolysis.

### Western blot and native gel assays

Equal amounts of purified proteasomes were resolved by SDS-PAGE, then transferred onto nitrocellulose membranes. Membranes were then probed for proteasome α-subunits, total Rpt6, and phospho-Rpt6. In-gel activity assays were performed by loading equal amounts of purified proteasomes onto native gradient gels (4-12%) and resolving overnight at 4°C. The gel was then soaked in developing buffer (50 mM Tris-HCl (pH 7.4), 5 mM MgCl_2_, 0.5 mM EDTA, 1 mM ATP) and supplemented with 50 µM suc-LLVY-AMC substrate for 30 minutes at room temperature. The gel was then imaged in a Protein Simple FluorChem E imaging system, resulting in fluorescent bands. Gels were then transferred to nitrocellulose and probed for 20S core subunits, Rpt6, and phospho-Rpt6 and then secondary HRP-conjugated antibodies. Resulting blots were digitized by scanning films and band intensities were quantified using NIH ImageJ.

### Slice Electrophysiology

Acute hippocampal slices were prepared from 3-8 week old mice with experimenter blind to genotype. Mice were anesthetized with isoflurane prior to decapitation and brain extraction into ice-cold sucrose-containing ACSF (in mM: 83 NaCl, 2.5 KCl, 1 NaH_2_PO_4_, 26.2 NaHCO_3_, 22 glucose, 72 sucrose, 0.5 CaCl_2_, 3.3 MgSO_4_). Tissue was sliced coronally into 350 µm slices using a Leica VT1200 vibratome. Slices were recovered in standard ACSF (in mM: 119 NaCl, 5 KCl, 1 NaH_2_PO_4_, 26 NaHCO_3_, 11 glucose, 2 CaCl_2_, 1 MgSO_4_) at 34°C for 30 minutes and at room temperature for at least 30 minutes prior to recordings. Slices were transferred to a submerged recording chamber and perfused with room-temperature (basal transmission experiments) or 30°C (potentiation experiments) oxygenated ACSF (with 100 µM picrotoxin).

A cluster stimulating electrode (FHC) was placed in the stratum radiatum of the CA1 region and current or voltage was injected using an ISO-Flex stimulus isolator (A.M.P.I.) triggered by a Clampex 10.3 (Molecular Devices) or custom IGOR Pro protocol. Recording electrodes were generated from thin-walled capillary tubing (Warner Instruments) using a horizontal pipette puller (P-97 Flaming/Brown Micropipette Puller, Sutter Instruments), resulting in a resistance of 1-3 MΩ, and pipettes were filled with ACSF. The recording electrode was placed 200-300 µm away from the stimulating electrode in the stratum radiatum, along a pathway parallel to the CA1 pyramidal layer. A second stimulating electrode was placed on the opposite side of the recording electrode to serve as a control pathway in LTP experiments. Field responses were recorded using an Axopatch 200B amplifier (Molecular Devices) and digitized using a Digidata 1322 digitizer (Molecular devices), and signals were acquired using Clampex 10.3 or IGOR Pro.

Field excitatory post-synaptic potentials (fEPSPs) were evoked using 100 µs pulses. Input-output relationships were determined by recording fEPSPs at a variety of stimulus intensities ranging from 30 µA to 130 µA and averaging five traces per intensity. Paired pulse facilitation was examined by generating pulses at variable separations (400, 200, 150, 100, and 50 ms) and taking the ratio of the second fEPSP amplitude to the first, with three recordings averaged for each separation. To study long-term potentiation (LTP), a stimulus intensity that produces a response of 0.2 – 0.4 mV was administered once every 15 seconds until a stable baseline of 15 minutes was reached. LTP was induced in one pathway using one set or four sets of 1-second 100 Hz trains separated by 20 seconds and post-LTP responses were recorded from both control and experimental pathways for 40-60 minutes after potentiation.

### Behavioral Assessment

#### Fear Conditioning

Mice were tested in individual conditioning chambers. The VideoFreeze system (Med Associates) was used to assess behavior as described previously (Anagnostaras et al., 2010; Carmack et al., 2014). Training consisted of one 10 min session. Mice were placed in chambers and baseline activity was assessed for 2 min. Mice then received 3 tone-shock pairings beginning at minutes 2, 3, and 4. Tone-shock pairings consisted of a 30-sec tone (2.8 kHz, 90 dB) that co-terminated with a 2-sec scrambled AC foot shock (0.75 mA, RMS). Twenty-four hours post-training, mice were returned to the training context. Once mice were placed in the chambers, freezing was measured for 5 minutes to assess fear to the context. Twenty-four hours after the context test a 5 min tone test was conducted to assess cued memory. The context was altered on multiple dimensions. White acrylic sheets were placed over the grid floors, a black plastic triangular insert was used to alter wall shape, and chambers were cleaned and scented with a 5% vinegar solution. Near-infrared light was used, in the absence of white light, to create a dark environment. The test consisted of a 2 min baseline period followed by the presentation of three 30 sec tones (2.8 kHz, 90 dBA) at minutes 2, 3, and 4. The tones matched those used during training.

Elevated Plus Maze: The plus maze (MED Associates) had two open and two enclosed arms (6.5 cm × 36 cm each) joined at a center hub (6.5 cm × 6.5 cm) elevated 74 cm from the ground. Testing lasted 5 min in dim light. The floor of the maze had near-infrared backlighting invisible to the mice to provide video contrast. Mice were tracked using a camera and video tracking software (Panlab Smart 3.0, Harvard Apparatus). Time spent in each arm and total distance traveled were recorded.

#### Spine density and immunohistochemistry

Adolescent (P21-P28) homozygous Rpt6 S120A and corresponding WT mice were given an anesthetic analgesic solution of ketamine and xylazine prior to intracardial perfusion with 0.9% saline, followed by 4% PFA fixative solution (20 mL 0.2 M PB, 10 mL ddH_2_O, 10 mL 16% PFA). Brains were postfixed for 1 hr, then 100 μm coronal sections were cut using a Vibratome. Penetrating microelectrodes were pulled from borosilicate capillary glass with filament (1 mm outer diameter/−0.58 mm inner diameter) and then backfilled with KCl (200 mM) and Alexa 594 hydrazide (10 mM) solution (Invitrogen). Using micromanipulation and visual guidance, CA1 neurons were filled via iontophoresis. Sections were postfixed for 10 min prior to mounting with Aqua Polymount (Polysciences Inc). Confocal microscopy was used to image secondary apical dendrites for analysis. The density, length, and width of dendritic protrusions were measured via Z-stacks in ImageJ using custom macros blind to genotype.

#### Preparation and transfection of organotypic slice cultures

Organotypic hippocampal slices were prepared from litter matched P6-P8 wild-type and S120A KI mice of both sexes, as previously described (Stoppini *et al.*, 1991). The cultures were transfected 1-2 d before imaging via biolistic gene transfer (180 psi), as previously described (Woods & Zito, 2008). 10-15 µg of EGFP (Clontech) was coated onto 6-8 mg of 1.6 µm gold beads.

#### Time-Lapse Imaging and Image Analysis for spine dynamics

EGFP-transfected CA1 pyramidal neurons [7-9 days in vitro (DIV)] at depths of 10-50 µm were imaged using a custom two-photon microscope (Woods *et al.*, 2011) controlled with ScanImage (Pologruto *et al.*, 2003). Slices were imaged at 29 °C in recirculating artificial cerebrospinal fluid (ACSF) containing in mM: 127 NaCl, 25 NaHCO_3_, 25 D-glucose, 2.5 KCl, 1.25 NaH_2_PO_4_, 1 MgCl_2_, and 2 CaCl_2_, aerated with 95%O_2_/5%CO_2_. For each neuron, image stacks (512 × 512 pixels; 0.035 μm/pixel) with 1 μm z-steps were collected from 5 to 6 segments of secondary dendrites (apical and basal) at 15 min intervals. To compare rates of activity-induced new spine formation in WT versus S120A KI mice, slices were treated with 30 µM bicuculline (Tocris, 1,000x aqueous stock) or vehicle. All shown images are maximum projections of three-dimensional (3D) image stacks after applying a median filter (3 × 3) to the raw image data. Spine formation rates were blindly analyzed in 3D using custom software in MATLAB.

#### Statistical Analyses

For comparison of two groups, we used a standard t-test. For groups of three or more, we performed a one-way ANOVA with Tukey’s post-hoc multiple comparison test or two-way ANOVA with Bonferroni’s post-hoc multiple comparison test. Significance threshold was set at p < 0.05. Behavioral data were analyzed using multivariate or univariate analyses of variance (ANOVAs). Significance was set at level *p* < 0.05 (SPSS Statistics Desktop, V22.0 or GraphPad Prism). Graphs depict Mean ± SEM.

## RESULTS

### Generation, purification, and analysis of Rpt6 S120A and S120D mutant proteasome composition and activity

In order to evaluate the functional relevance of Rpt6 S120 phosphorylation in the brain and at synapses, we generated two novel mouse lines by mutating Rpt6 S120 to either alanine (S120A) which blocks phosphorylation, or to aspartate (S120D), which mimics phosphorylation due to its negatively charged carboxyl group (Supplementary Figure 1A-C). By visual inspection, Rpt6 S120A and Rpt6 S120D homozygous mutants displayed normal body size, feeding, and mating behaviors. In addition, there were no obvious gross brain anatomical differences (Supplementary Figure 1D).

We evaluated proteasome peptidase and ATPase activity of proteasomes purified from Rpt6 WT and mutant mice. The gentle affinity purification protocol we utilized more efficiently maintains interactions between the regulatory and catalytic particles of the proteasome, as well as those between the proteasome and its interacting proteins (Besche & Goldberg, 2012). In order to verify efficient purification of WT and mutant proteasomes, we resolved the purification products on SDS-PAGE gels. Western blots showed high concentrations of 20S subunits as well as total Rpt6. When we probed for phospho-Rpt6 using our affinity purified phospho-Rpt6 antibody, S120A mutants showed little to no cross-reactivity, while S120D showed marginal cross-reactivity, as expected (Figure 1A). In order to evaluate composition and functionality of our purified proteasomes, we ran the purification products on a native gel and performed an in-gel fluorescence assay utilizing the LLVY-AMC substrate. After electrophoresis, we bathed the gel in a buffer containing the substrate and imaged it on a Protein Simple UV imager, rendering bands for both single and doubly capped proteasomes (Figure 1B), verifying that our proteasomes were intact and fully functional following purification. We further confirmed the composition of purified proteasomes by mass spectrometry analysis (data not shown).

**Figure 1:**
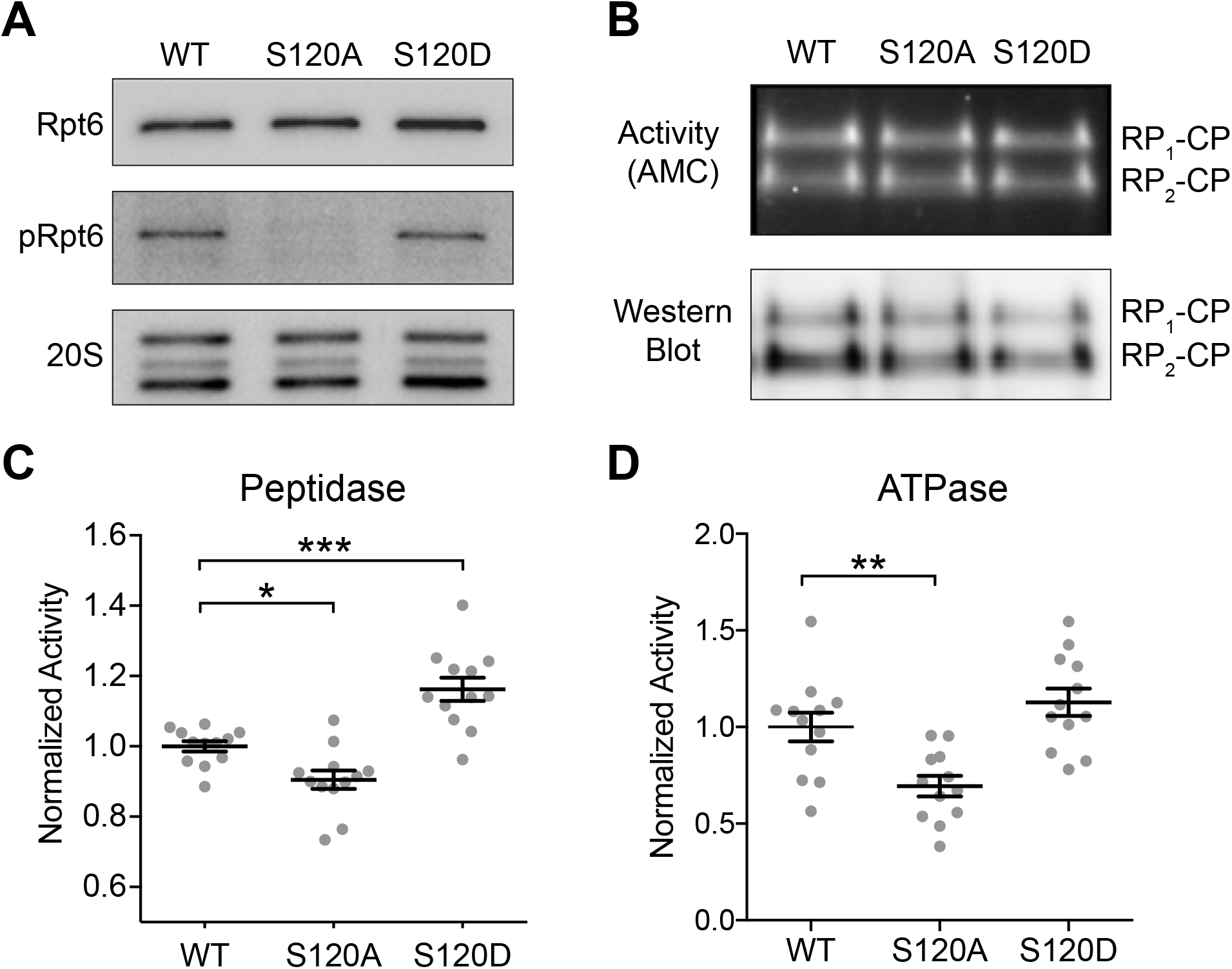
Proteasome protein degradation and ATP hydrolysis kinetics are significantly altered in Rpt6 S120A and S120D knock-in mice. **(A)** Representative western blot analysis of proteasomes purified using Ubl/UIM method, and verification of phospho-Rpt6 antibody specificity to WT Rpt6 with expected cross reaction with S120D Rpt6 (performed for each purification). **(B)** Native gel fluorescent activity assay and tandem western blot with two bands representing singly and doubly capped proteasomes (performed for each purification). **(C)** Fluorescent peptidase activity assay; S120A has significantly decreased activity (p < 0.04, post-hoc Tukey, n = 12), S120D shows increased activity (p < 0.001, post-hoc Tukey, n = 12), and ANOVA indicates significant difference within the group (F = 25.26, p < 0.001). **(D)** Malachite green ATPase assay; S120A displays significantly lower activity (p = 0.007, post-hoc Tukey, n = 12), while S120D shows increased activity (p = 0.37, post-hoc Tukey, n = 12), and the one-way ANOVA indicates significant difference within the group (F = 11.21, p < 0.001).

In order to quantitatively evaluate proteasome activity, we utilized kinetics assays to measure peptidase and ATPase activities to determine if they exhibit the same differences as they did in cultures. We found that Rpt6 S120A phospho-dead mutants showed significantly reduced peptidase activity as compared to WT (p < 0.04; n = 12), while Rpt6 S120D phospho-mimetic mutants exhibit significantly increased peptidase activity (p < 0.001; n = 12) (Figure 1C).

Rpt6 is one of six ATPase subunits in the 19S regulatory particle. These subunits hydrolyze ATP, providing the mechanical energy necessary to unfold target substrates and move them inside the catalytic core (Yedidi *et al.*, 2017). We therefore investigated the ATPase activities of our purified wild type and mutant proteasomes. Indeed, we found that the S120A mutant displayed a reduced rate of ATPase activity compared to wild type (p = 0.007; n = 12), and the S120D mutant showed a slight but statistically insignificant increase in activity (p = 0.37; n = 12) (Figure 1D).

### Elimination of phosphorylation dynamics do not effect basal synaptic transmission nor LTP paradigms of synaptic plasticity

We first assessed whether Rpt6 mutations affect basal synaptic transmission, using Schaffer collateral (CA3-CA1) stimulation in acute hippocampal slices from mutant and wildtype mice. We observed that slices from both S120A and S120D animals display a normal input-output relationship when compared to slices obtained from age-matched wild-type mice (Figure 2A, B). This indicates that the hippocampal circuit has developed normally and has roughly the same amount of synaptic connectivity as a wild-type hippocampus. Thus, these specific mutations to S120 do not obviously impact the formation and maintenance of excitatory synapses in the hippocampus, and do not appear to strongly impact synaptic strength at these synapses. To assess whether presynaptic release mechanisms were altered in S120A and S120D mice, we next utilized a paired pulse facilitation assay. By stimulating Schaffer collateral axons twice at varying pulse separations, we revealed strong facilitation of fEPSPs in wild-type slices at small separations (50 ms, 100 ms). S120A and S120D slices displayed identical levels of facilitation, indicating that presynaptic neurotransmitter release is intact and has normal responses to increased presynaptic calcium (Figure 2C, D).

**Figure 2:**
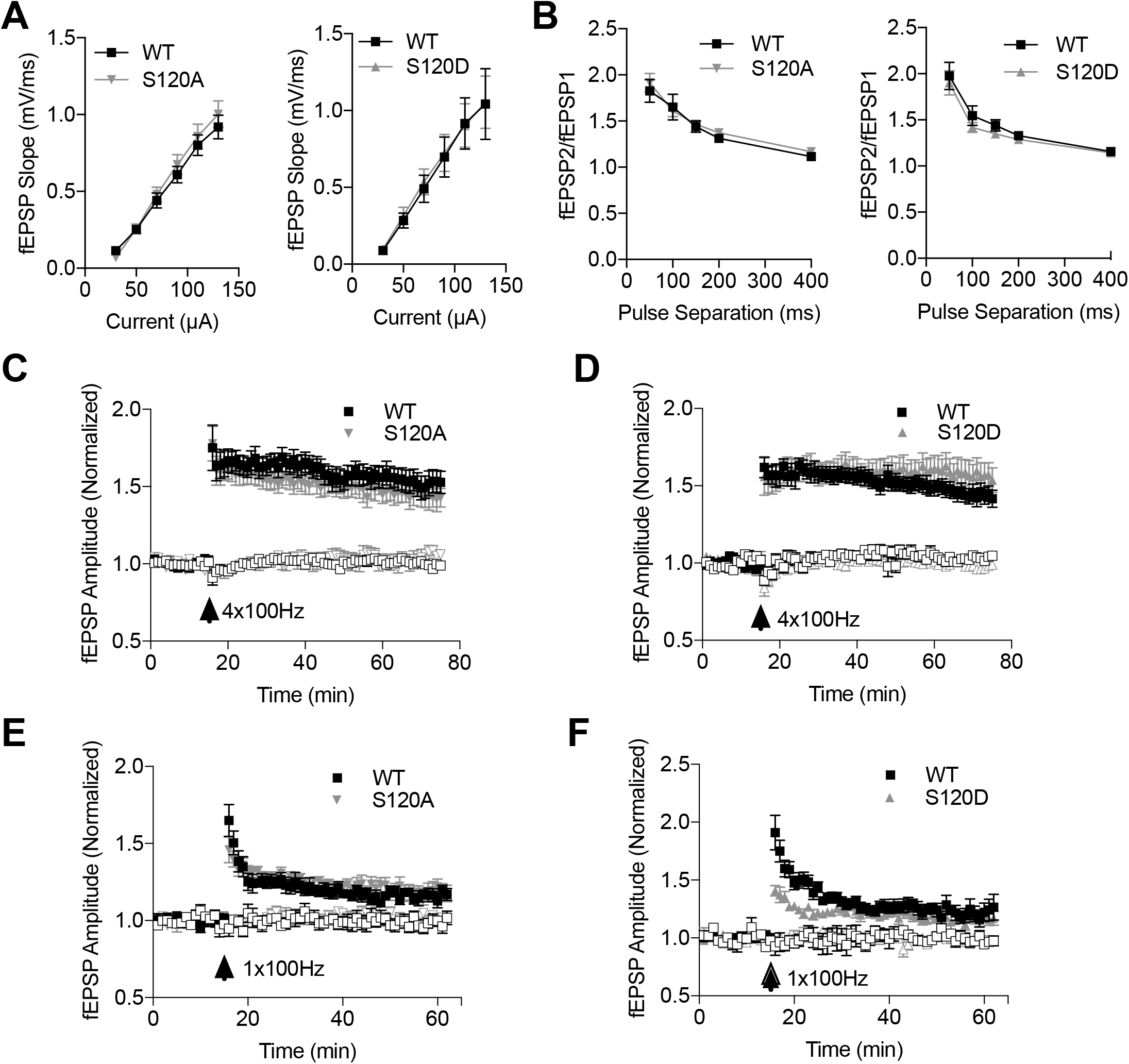
Basal synaptic transmission and LTP are unaltered in Rpt6 S120A and S120D knock-in mice. **(A)** Input/output curve in slices from wildtype and S120A mice (n = 15 slices per genotype), depicting increasing field EPSP responses to increased amplitude of Schaffer collateral stimulation. **(B)** Input/output curve in acute hippocampal slices from wildtype and S120D mice (n = 11 slices per genotype). **(C)** Two successive stimuli with short separation delivered to the Schaffer collateral of acute hippocampal slices leads to enhancement of fEPSP amplitude (paired pulse facilitation) in wildtype and S120A mice (n = 16 slices per genotype). **(D)** Paired pulse facilitation in wildtype and S120D mice (n = 10 slices). **(E)** Delivery of four 1-second trains (20 s apart) of 100 Hz stimulation causes potentiation of fEPSP amplitude in acute hippocampal slices from wildtype and S120A mice (n = 6, 9 slices, respectively). **(F)** LTP induction in response to four 1-second 100 Hz trains in wildtype and S120D mice (n = 9, 10 slices, respectively). **(G)** LTP induction in response to a single 1-second 100 Hz train in slices from wildtype and S120A animals (n = 6, 7 slices, respectively). **(H)** LTP induction in response to a single 1-second 100 Hz train in wildtype and S120D animals (n = 5, 6 slices, respectively).

Since previous work with proteasome inhibitors pointed to an important role for these complexes in mediating long-term potentiation (LTP), we next sought to determine whether our Rpt6 mutations would affect this plasticity paradigm. We generated acute hippocampal slices from young (P21-P27) age-matched mice and induced LTP in the CA3-CA1 pathway using four trains of 1-second 100 Hz stimulation. We observed that hippocampal slices from S120A and S120D animals undergo normal levels of fEPSP potentiation compared to wild-type mice (Figure 2E, F). Due to previous studies suggesting that proteasome activity can modulate LTP induction thresholds (Dong et al., 2008), we next used a weaker stimulation paradigm, instead stimulating with a single 1-second 100 Hz train. While both mutants displayed normal levels of slight potentiation 15-30 minutes after LTP induction (Figure 2G, H), we did observe a suppression of post-tetanic potentiation and early LTP in S120D animals (Figure 2G). However, since potentiation eventually reached wild-type levels, it appears that neither mutation has a strong effect on LTP.

### Spine density and outgrowth dynamics are not altered in S120A or S120D knock-in mice

Proteasome function is critical in the maintenance of synaptic plasticity and spine outgrowth. We previously showed that proteasome function is necessary for the formation of new spines in response to increased neuronal activity (Hamilton *et al.*, 2012). Furthermore, we have shown that increasing synaptic activity also augments proteasome peptidase and ATPase activity. Therefore, we wanted to investigate if the Rpt6 S120A knock-in mutant that cannot be phosphorylated had any effect on the activity-induced formation of new spines. Hippocampal pyramidal neurons in organotypic slice cultures of WT and litter-matched homozygous S120A mutants were transfected with EGFP and imaged over time using a two-photon microscope. Dendrites of EGFP-expressing CA1 neurons were imaged at 15 min intervals before and after being treated with either vehicle, or 30 µM bicuculline (Figure 3A). Spines were counted and compared to their vehicle controls. We found no difference in the basal spine density measured before treatment (p = 0.839, n = 6 for WT and S120A). We also observed no significant changes in the overall formation of new spines under basal (veh-treated) conditions or following increased activity (bic-treated) (p = 0.891, n = 6 for both WT and S120A), and bicuculline successfully induced significant spine outgrowth in both genotypes (p = 0.02 and p = 0.04 for WT and S120A, respectively) (Figure 3B-C). To further support this finding, we prepared sections of fixed brains from adolescent WT and S120A mice, micro-injected CA1 neurons with Alexa Fluor 594 fluorescent dye, and counted the individual number of spine heads on 20 micron segments of secondary dendritic branches using confocal microscopy (Supplemental Figure 2A-B). We found that there was no statistically significant difference between the WT control and S120A mice (p = 0.828, n = 56 for WT and n = 51 for S120A), indicating that dendritic spines in the hippocampus develop normally in the absence of S120 phosphorylation dynamics.

**Figure 3:**
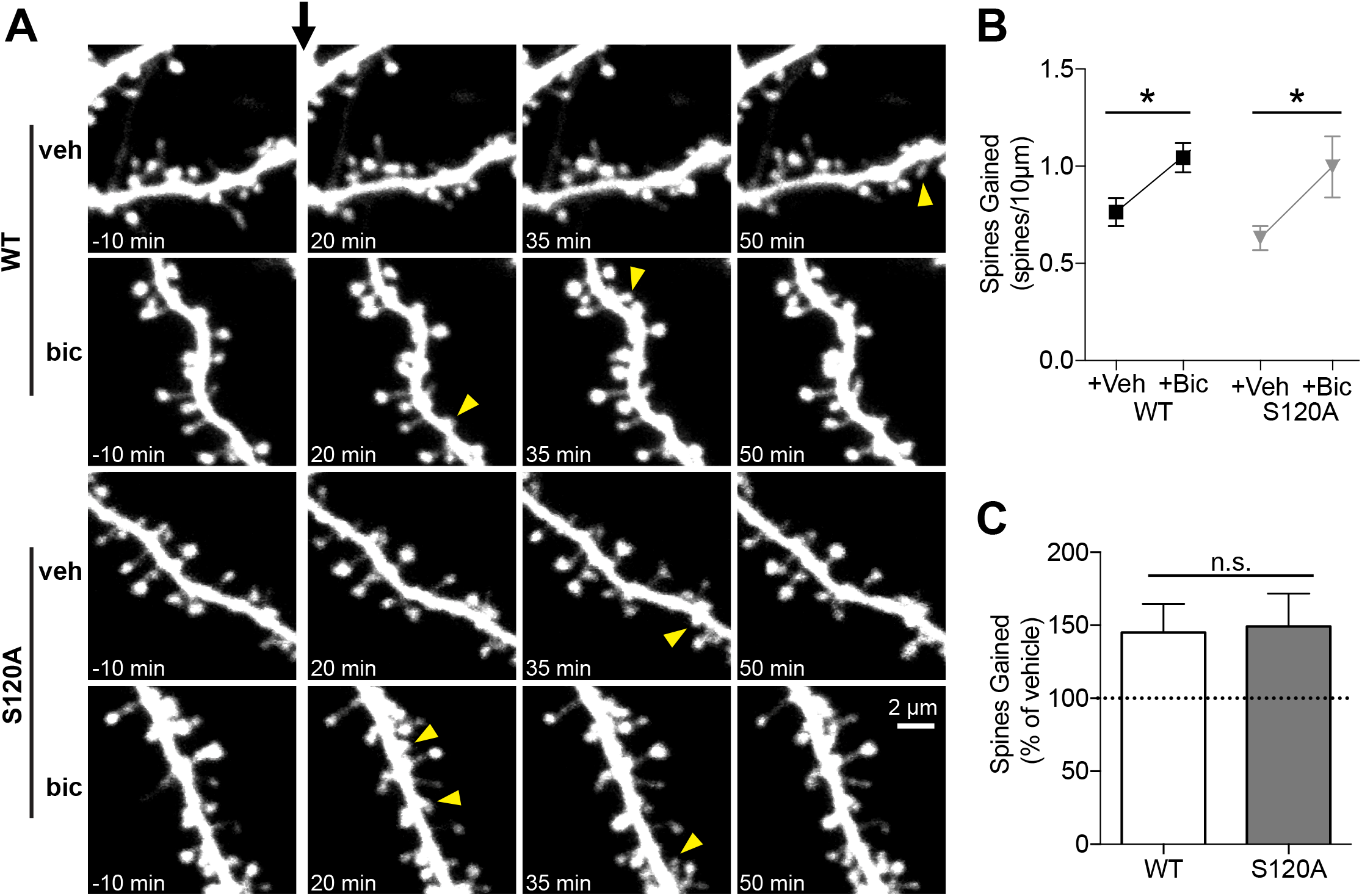
Neither basal spine density nor outgrowth dynamics are altered in S120A knock-in mice. **(A)** Images of dendrites from EGFP-expressing CA1 pyramidal neurons from WT and S120A animals at 7-9 DIV before and after the addition of vehicle or bicuculline (30 μM) at t = 0 (black arrow). Yellow arrowheads indicate new spines. **(B)** Quantification of number of spines gained per 10 µm with addition of vehicle or bicuculline, in WT (n = 6, 6, respectively) and S120A (n = 7, 6, respectively). Bicuculline induced significantly more spine outgrowth than vehicle in both WT and S120A neurons (p = 0.02 and p = 0.04, respectively), but no significant difference was observed in bic-induced outgrowth between WT and S120A (p = 0.79 with t-test; ANOVA of interaction, F(1,21) = 0.197, p = 0.66). **(C)** Quantification of normalized spine gain (% of vehicle) in neurons from WT and S120A slices, illustrating similar bic-induced spine outgrowth in both genotypes (p = 0.89 with t-test).

### Learning and memory are unaltered in Rpt6 S120A and S120D knock-in mice

We next sought to evaluate the impact of our knock-in mutations on learning and memory, due to previous studies linking S120 phosphorylation by CaMKII to long-term memory formation (Jarome et al., 2013). First, to ensure that our mutations did not induce any abnormal anxiety-related behaviors or locomotor deficits that may affect our interpretation of other behavioral data, we assessed these mutants using an elevated plus maze paradigm. S120A (n = 8) and WT (n = 9) mice did not show any difference in total distance traveled during this test (p = 0.16), nor did S120D (n = 14) and their WT controls (n = 13; p = 0.80) (Supplementary Figure 3A-B). Percentage of time spent in the open and closed arms did not differ between WT and S120A mice (p = 0.82 and p = 0.08, open and closed respectively), nor did it differ between WT and S120D mice (p = 0.17 and p = 0.30, open and closed respectively) (Supplementary Figure 3C-D). Thus, these knock-in mice show normal behavior overall and do not display any apparent anxiety-related phenotypes.

The effects of blocking and mimicking Rpt6 phosphorylation at serine 120 on long-term associative memory were examined using Pavlovian cued and contextual fear conditioning. Baseline activity, measured during the first 2 minutes of training, did not differ between S120A (n = 16) and WT (n = 8) mice [F(1,22) = 0.25, p = 0.62] (Figure 4A). However, shock reactivity in S120A mice was slightly dampened [F(1,22) = 4.33, p = 0.049] (Figure 4A). The difference in shock reactivity did not seem to influence memory for the task. When assessed 24 hours later, we did not observe any differences in cued [F(1,22) = 0.119, p = 0.73] nor contextual [F(1,22) = 1.645, p = 0.21] memory (Figure 4C-D). Similarly, S120D (n = 21) and WT (n = 13) mice displayed no difference in baseline activity [F(1,32) = 1.25, p = 0.27], but we observed a reduction in shock reactivity [F(1,32) = 10.48, p < 0.005] (Figure 4B). However, this difference once again did not affect learning and memory in this task, as both cued [F(1,32) = 1.20, p = 0.28] and contextual [F(1,32) = 0.087, p = 0.77] fear memory were intact in the knock-in mice (Figure 4C-D).

**Figure 4:**
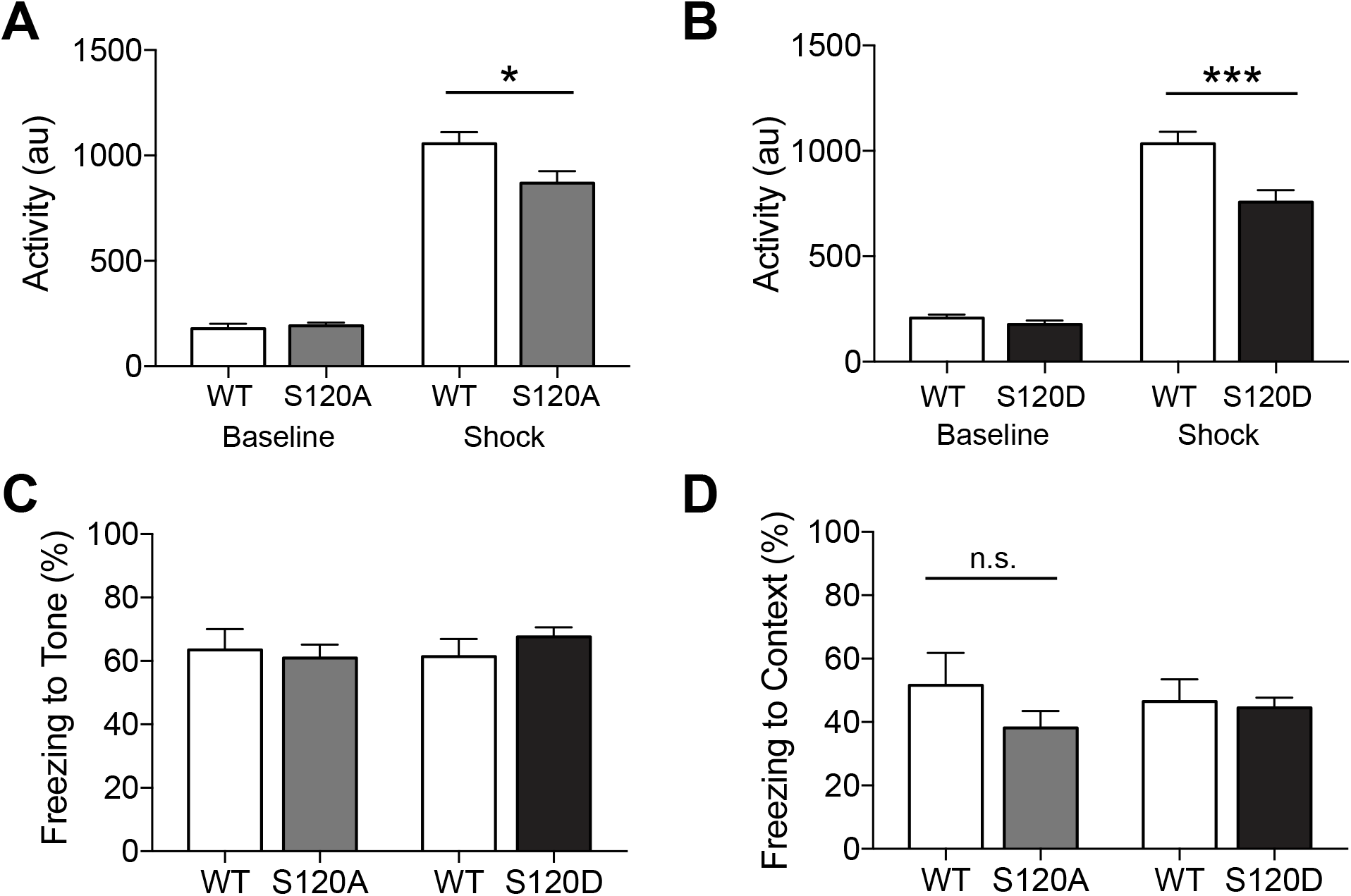
Contextual and cued fear conditioning are unaltered in Rpt6 S120A and S120D knock-in mice. **(A)** Activity during 2-minute baseline and activity during 2-second shock in WT (n = 8) and S120A (n = 16) mice, demonstrating normal baseline activity (p = 0.62) but slightly attenuated shock reactivity (ANOVA, F(1,22) = 4.33, p < 0.05) in S120A mice. **(B)** Same as **A**, but for WT (n = 13) and S120D (n = 21), depicting similar baseline activity (p = 0.27) but reduced shock reactivity (ANOVA, F(1,32) = 10.48, p < 0.005). **(C)** Percent time spent freezing to three sequential tone presentations in a novel context, during post-training (24 h) test; S120A and S120D knock-in mice show similar amounts of freezing compared to respective WT mice (p = 0.21, p = 0.28, respectively). **(D)** Percent time spent freezing to initial training context during 5-minute test, 24 h after training; S120A and S120D mice show normal amounts of freezing (p = 0.73, p = 0.77, respectively).

## DISCUSSION

In an effort to further understand the importance of proteasome phosphorylation and its functional relevance at synapses, we created novel mutant mouse lines which constitutively block or mimic phosphorylation of Rpt6 at ser-120. The impetus to create these mice stemmed from several bodies of work from our and other laboratories which describe how the 26S proteasome is regulated in neurons by synaptic activity. In 2006, Bingol and Schuman provided the first evidence that the 26S proteasome may play an instructive role at synapses in work that described the activity- and NMDA receptor-dependent translocation and sequestration of proteasomes at synapses (Bingol and Schuman, 2006). We then showed that neuronal activity regulates proteasome activity by a mechanism involving CaMKIIα-mediated phosphorylation of Rpt6 at ser-120 (Djakovic et al., 2009; 2012). Subsequently, it was shown that autophosphorylated CaMKIIα acts as a scaffold for proteasomes in dendritic spines (Bingol et al., 2010). In further investigations, we found that homeostatic scaling of synaptic strength induced by chronic application of bicuculline or tetrodotoxin is both mimicked and occluded by altered Rpt6 phosphorylation (Djakovic *et al.*, 2012). Furthermore, it was demonstrated that the acute inhibition of the proteasome or expression of Rpt6 S120A phospho-dead mutant blocks activity-dependent new spine generation (Hamilton *et al.*, 2012). Collectively, these data suggested that CaMKIIα-dependent phosphorylation of Rpt6 at S120 may be an important regulatory mechanism for proteasome-dependent remodeling of synapses.

We first evaluated proteasome peptidase and ATPase activity of intact 26S proteasomes purified from Rpt6 WT, S120A, and S120D mutant mice. Compared to proteasomes purified from WT mice, we found peptidase activity and ATPase activity of proteasomes purified from Rpt6 S120A and S120D mutant to be decreased and increased, respectively, corroborating our previous findings with ectopically expressed Rpt6 S120A and S120D mutants (Djakovic *et al.*, 2009). Unlike proteasome inhibitors, these mutations only alter the activities of the proteasome by approximately 15 to 20%. To determine the functional relevance of these mutations on synapses we evaluated basal synaptic transmission, LTP, spine density and new spine growth, and fear memory. Unlike the striking differences we observed when Rpt6 S120A or S120D mutants are ectopically expressed in neurons (Djakovic *et al.*, 2012; Hamilton *et al.*, 2012), we observed no significant differences in the Rpt6 S120A and S120D knock-in mice.

Previous work suggested an important role for proteasome-dependent protein degradation in learning and memory. Early studies observed impairments in one-trial avoidance learning, spatial learning, and aversive taste memory formation upon treatment with proteasome inhibitors (Lopez-Salon et al., 2001; Artinian et al., 2008; Rodriguez-Ortiz et al., 2011). Supporting a specific role for S120 phosphorylation in learning, it was more recently observed that fear conditioning transiently increases phosphorylation by CaMKII to stimulate proteasome activity (Jarome et al., 2013), and that this CaMKII-based mechanism of proteasomal regulation may be essential for memory reconsolidation (Jarome et al., 2016). In light of this, the lack of behavioral impairments in our S120 knock-in mice is somewhat surprising. This either suggests that this particular site is not as essential as previously thought, or that alternate mechanisms are able to differently regulate proteasomal activity to achieve the same end result.

While numerous mechanisms of regulation of the proteasome and its function have been discovered and well characterized, our results demonstrate that Rpt6 S120 phosphorylation dynamics may not play essential roles in the brain. This could be due to an ability of neurons to compensate for minor alterations in proteasome function using either alternative mechanisms of protein degradation, such as lysosomal degradation, or by simply changing transcription/translation rates. Recent work has illustrated the interplay and feedback loops between the UPS and the lysosome system, often showing that when one pathway is inhibited, the other is upregulated in order to compensate for the disturbance. Specifically, activation of mTORC1, which inhibits lysosomal degradation, has been shown to increase the number of intact and active proteasomes through the activation of NRF1 (Zhang *et al.*, 2014; Zhang & Manning, 2015). Additionally, it has been shown that certain transcription factors for proteasomal subunits are also substrates of the proteasome, creating a feedback loop where inhibition increases the transcription/translation of proteasome subunits, leading to an increase in active holoenzymes (Xie & Varshavsky, 2001). Interestingly, some of these compensation mechanisms are only observed under low concentrations of proteasome inhibitors, and are absent under high levels of inhibition (Sha & Goldberg, 2014), indicating specialized compensatory mechanisms for minor alterations in activity.

While several other proteasome subunits have been shown to be phosphorylated (Wang *et al.*, 2007; Murata *et al.*, 2009; Tomko & Hochstrasser, 2013; Schmidt & Finley, 2014), the functional relevance of these modifications on proteasome function remains poorly understood. Recently, the cell cycle-dependent phosphorylation of Rpt3 at Thr25 was shown to be important for cell cycle progression and tumorigenesis (Guo *et al.*, 2016), which supports the idea that single site phosphorylation events are clearly important and perhaps rate-limiting on specific proteasome-dependent biological processes.

Though altered Rpt6 S120 phosphorylation dynamics did not affect basal synaptic transmission, LTP, spine dynamics, and fear memory, this does not rule out that there exist biological pathways where downstream effects are dependent on small changes in proteasome activity directly linked to phosphorylation of Rpt6. For instance, our laboratory has recently discovered that behavioral sensitization to cocaine is highly dependent on Rpt6 phosphorylation by CaMKIIα after cocaine administration. In that study, we found that cocaine increases CaMKIIα-dependent phosphorylation of Rpt6, and concomitantly increases proteasome activity in the nucleus accumbens and in the pre-frontal cortex. In contrast, cocaine does not increase proteasome activity in Rpt6 S120D mutants, and strikingly, we found a complete absence of cocaine-induced locomotor sensitization in the Rpt6 S120D mice (Gonzales *et al.*, 2017), which suggests a critical role for Rpt6 S120 phosphorylation and proteasome function in the regulation cocaine-induced behavioral plasticity. Furthermore, blocking phosphorylation of Rpt6 at S120 appears to sensitize both yeast and mice to pathogenic effects of Huntington protein aggregation, suggesting that Rpt6 S120 phosphorylation might be engaged under various forms of stress (Lin *et al.*, 2013; Marquez-Lona *et al.*, 2017). It is therefore plausible that Rpt6 phosphorylation could be important and perhaps rate-limiting for proteasome function in other biological pathways in the CNS yet to be identified, or perhaps in specific cell types under conditions that require a high degree of rapid protein turnover.

---

**Supplemental Figure 1:**
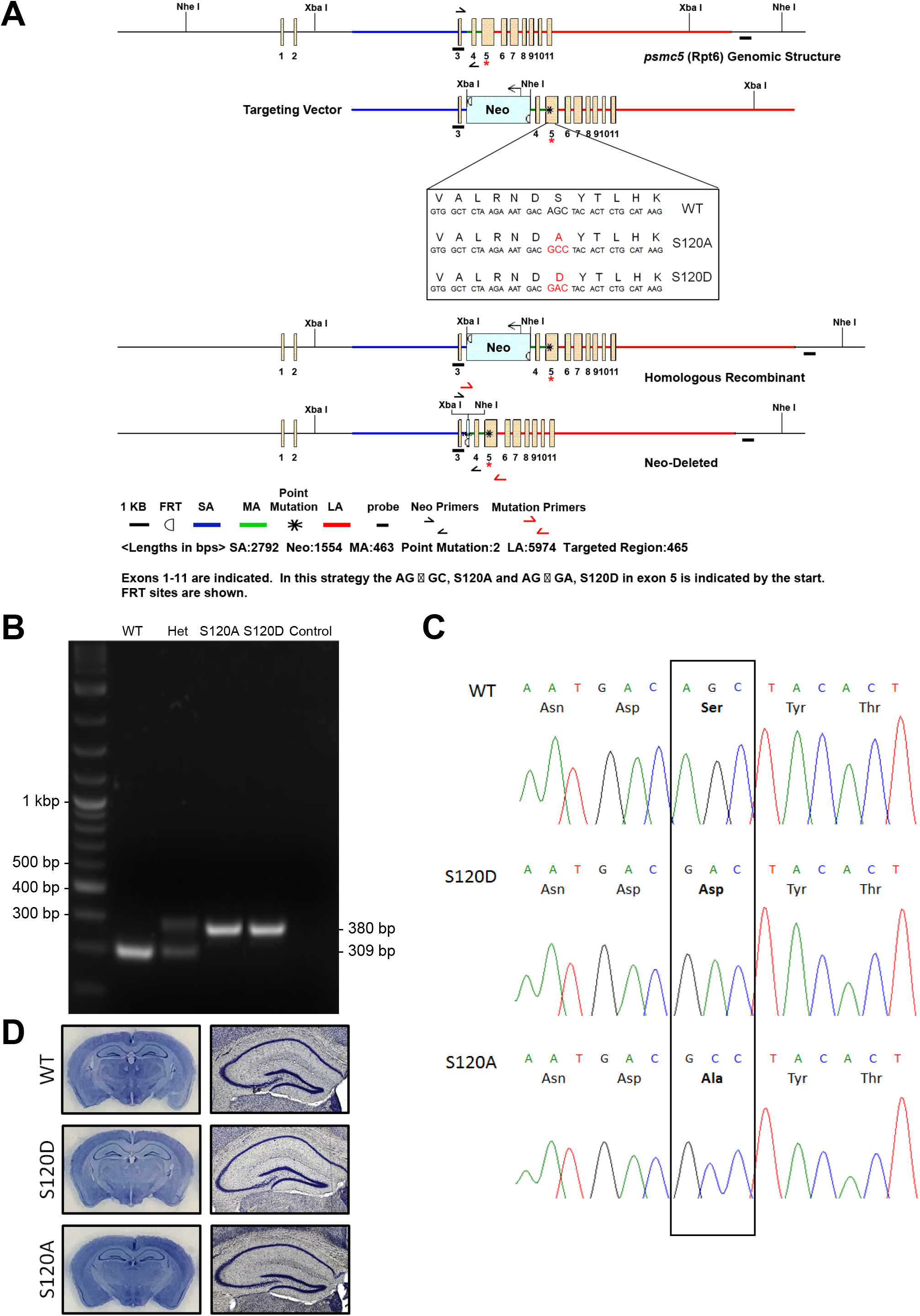
Generation of *Rpt6* S120A and S120D Knock-in (KI) mice. **(A)** Schematic of targeting strategy. Genomic DNA structure of *psmc5 (Rpt6*) region is shown. The targeting vector was designed such that the long homology arm (LA) extends 5.98 kb 3’ to the first point mutation (asterisk; AG → GC, S120A; AG → GA, S120D) in exon 5. The FRT flanked Neo resistance cassette was inserted 463 bp 5’ to the point mutation. The short homology arm (SA) extends 2.79 kb 5’ to the FRT flanked Neo cassette. **(B)** Tail genomic DNA was analyzed by PCR screening for genotyping and to verify deletion of the Neo cassette. **(C)** Electropherograms confirming the presence of the mutation in homozygous Rpt6 S120A and S120D mutant male mice. **(D)** Representative images of Nissl-stained fixed whole brain coronal sections of 60 day old mice.

**Supplemental Figure 2:**
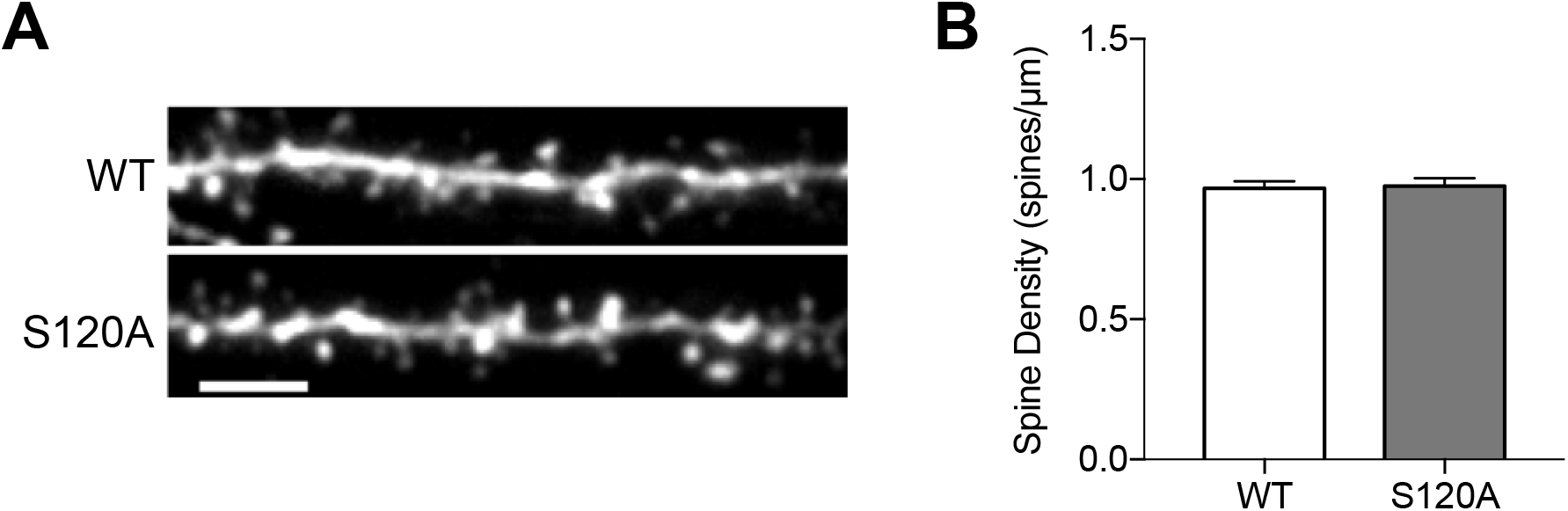
Spine density is not altered in S120A knock-in mice. **(A)** Representative proximal dendritic segments from hippocampal pyramidal neurons in fixed slices obtained from WT and homozygous S120A mutant littermates, after filling by targeted microinjection of Alexa Fluor 594 Hydrazide. **(B)** Basal spine density quantified by counting the individual number of spine heads on 20 micron segments of secondary dendritic branches in 63X confocal stacks (WT: n = 56 segments from 12 cells; S120A: n = 51 segments from 10 cells), showing no significant difference in density (p = 0.83).

**Supplemental Figure 3:**
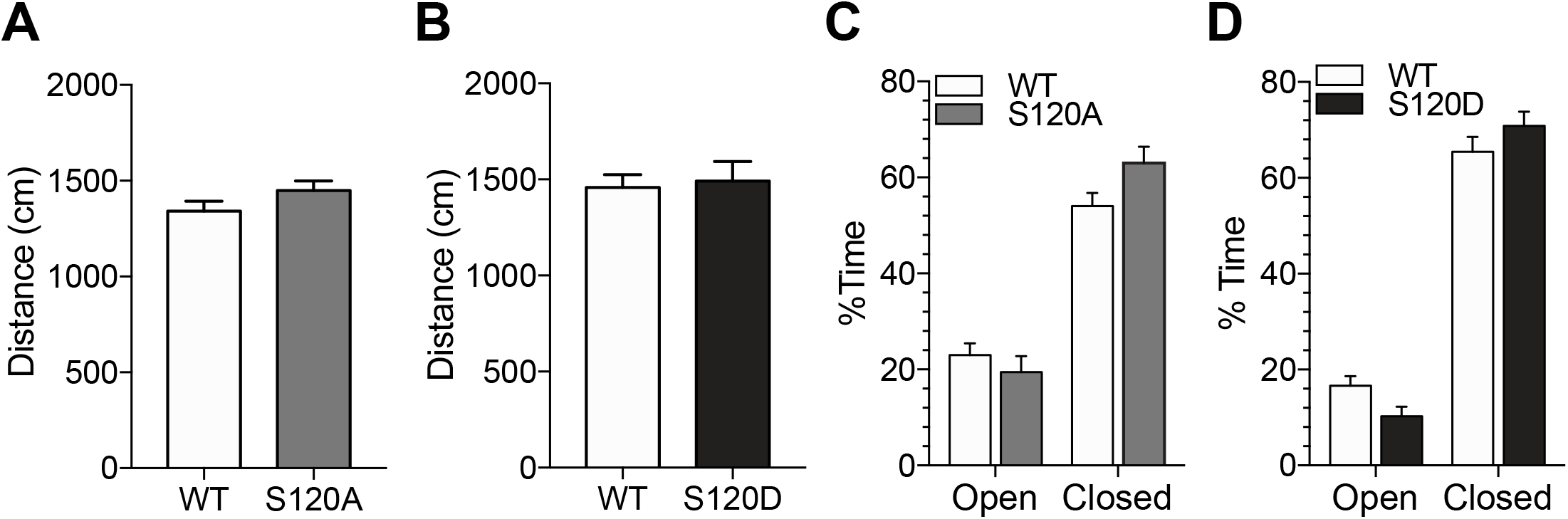
Performance on the elevated plus maze was not impaired in S120 knock-in mice. **(A)** Total distance travelled during elevated plus maze assay, demonstrating no differences between WT (n = 9) and S120A (n = 8) mice (p = 0.16, t-test). **(B)** Same as **A**, for WT (n = 13) and S120D (n = 14) (p = 0.80, t-test). **(C)** Percent time spent in each arm of the elevated plus maze (open vs closed) did not differ between S120A (n = 8) and WT (n = 9) mice (p = 0.82 and p = 0.08, open and closed respectively, post-hoc Bonferroni). (D) Same as **C**, for S120D (n = 14) and WT (n = 13) (p = 0.17 and p = 0.30, open and closed respectively, post-hoc Bonferroni).

